# A core microbiota of the plant-earthworm interaction conserved across soils

**DOI:** 10.1101/571240

**Authors:** Samuel Jacquiod, Ruben Puga-Freitas, Aymé Spor, Arnaud Mounier, Cécile Monard, Christophe Mougel, Laurent Philippot, Manuel Blouin

## Abstract

Microorganisms participate in most crucial soil functions and services benefiting human activities, such as biogeochemical cycles, bioremediation and food production. Their activity happens essentially in hotspots created by major soil macroorganisms, like rhizosphere and cast shaped by plants and earthworms respectively^1^. While effects of individual macroorganism on soil microbes are documented, no studies attempted to decipher how the mosaic of microhabitats built by multiple macroorganisms and their interaction determine the structure of microbial communities. Here we show a joint shaping of soil bacterial communities by these two macroorganisms, with a prevalent role of plants over earthworms. In a controlled microcosm experiment with three contrasted soils and meticulous microhabitat sampling, we found that the simultaneous presence of barley and endogeic earthworms resulted in non-additive effects on cast and rhizosphere bacterial communities. Using a source-sink approach derived from the meta-community theory^2,3^, we found specific cast and rhizosphere *core microbiota*^4,5^ of the plant-eartworm interaction, detected in all soils only when both macroorganisms are present. We also evidenced a *core network* of the plant-earthworm interaction, with cosmopolitan OTUs correlated both in cast and rhizosphere of all soils. Our study provides a new framework to explore aboveground-belowground interactions through the prism of microbial communities. This multiple-macroorganisms shaping of bacterial communities also affects fungi and archaea, while being strongly influenced by soil type. Further functional investigations are needed to understand how these *core microbiota* and *core network* contribute to the modulation of plant adaptive response to local abiotic and biotic conditions.

## Introduction

While the structuring effect of plants on soil microbial communities is well-documented, the overlooked role of earthworms on their abundance and activity may be as important^6,7^, given that they represent the highest animal biomass in soils^8^. Plants and earthworms are the major soil ecosystem engineers^9^, shaping microhabitats^10^ populated with microbes originating either from endogenic (e.g. endophytes and earthworm guts) or environmental (e.g. bulk soil) sources, namely the rhizosphere and drilosphere (casts and burrows)^1^. As such, plants and earthworms may be regarded as competing biotic entities for the steering of soil microbial communities and functions. Earthworms may even affect the assembly outcome of rhizosphere microbial communities^11^, but the reciprocal action of plant on drilosphere microbiota has never been investigated. Nevertheless, positive interaction between these two macroorganisms is the rule, as earthworms increase plant growth by ~25%^12,13^. Furthermore, converging observations suggest that part of the mechanics governing plant-earthworm relationships may pass through microbes^14,15^. Therefore, they may also be regarded as partners in shaping soil microbial communities.

Building on the concept of *core microbiota*^4,5^ associated with a host, we questioned the existence of such an extended entity resulting from the interaction of multiple hosts by focusing on plant and earthworm presence across soils. By looking at the right scale through the prism of microhabitats (e.g. rhizosphere, cast and bulk), we expected to capture the reciprocal influences of plant and earthworm on microbial communities, be they competitive, collaborative or neutral. We looked at this interaction in three contrasted soils (clayed, loamy, sandy) using microcosms containing either: i) a plant (barley, *Hordeum vulgare*), ii) endogeic earthworms (*Aporrectodea caliginosa*), iii) both macroorganisms and iv) a control without macroorganim. After one month, we applied a meticulous soil dismantling of each microcosm to separately harvest soil matrix (no macroorganisms), rhizosphere, cast and their respective bulk soils (Supporting File 1). We focussed on bacteria through 16S rRNA gene amplicon sequencing, because of their importance in the plant-earthworm interaction^16^. We also monitored the total abundance of fungi, archaea and bacteria using real-time quantitative PCR, as well as several plant traits.

## Results and discussion

In accordance with the literature^12,13^, we observed a positive effect of earthworms on shoot biomass whatever the soil (+21%), associated with an increase in height (+5%) and leaf surface area (+11%) in clayed and sandy soils (Tab.1). Beta-diversity analysis of bacterial community profiles revealed a strong soil effect (Supporting File 2 and 3), calling for a refined analysis *by soil* (Fig.1). Rhizosphere communities differed from those of other microhabitats, revealing the systematic plant influence whatever the soil (CAP1, 18-24%) while earthworms effect was weaker and soil-dependent (CAP2, 4-8%). Bulk and cast communities in earthworms treatment clustered away from the control bulk matrix in the clayed soil, indicating an overruling earthworm effect (yellow apart from blue, Fig.1a). However, earthworms had a weaker effect in the sandy soil, as bacterial communities in bulk and cast were similar to the control (yellow close to blue, Fig.1c). The loamy soil had an intermediate profile (Fig.1b).

**Fig.1:**
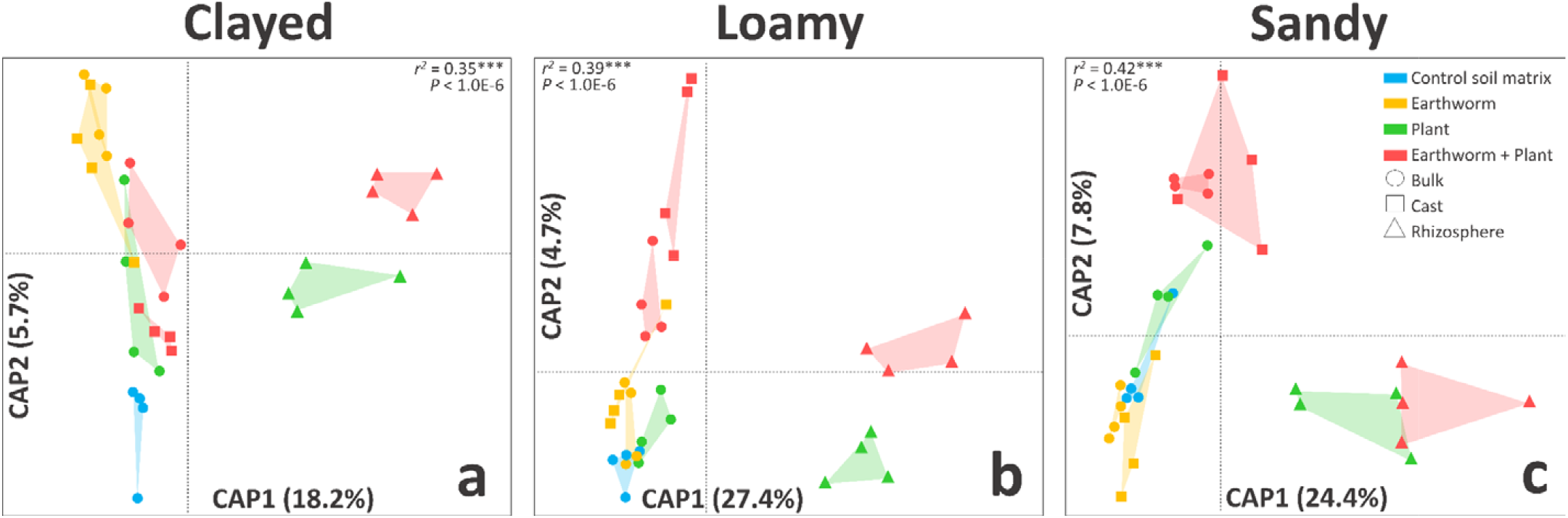
Canonical analysis of principal coordinates of bacterial communities in each soil (CAP, variance adjusted weighted unifrac distances). CAP1 and 2 represent the constrained components with their respective percentage of the total variance explained. Each model was validated using 10.000 permutations. The four treatments are indicated by different colors, while microhabitats are indicated by different marker shapes (Total n = 96).

**Tab. 1:** Effect of earthworms presence on barley traits at harvest in each soil. Three traits were measured (rows), including height (the longest leaf length, which was always the highest in our case), dry shoot weight and leaf surface area. Statistical significance was tested using two-sided, two-sample Student tests (*p* < 0.05) to compare average values (± standard error of the mean) between the condition without (w/o ew) and with (w ew) earthworms. Lowercase letters indicate statisticaly significant difference between tested average values (“a”: highest, “b”: lowest). All tested conditions were set with five biological replicates (n = 60).

| Soil | clayed |  | loamy |  | sandy |  |
| --- | --- | --- | --- | --- | --- | --- |
| Earthworms | - | + | - | + | - | + |
| Height (mm) | 34.80±0.26<sup>b</sup> | 37.10±0.80<sup>a</sup> | 30.30±0.51 | 31.10±0.60 | 32.10±0.56<sup>b</sup> | 34.30±0.44<sup>a</sup> |
| Dry shoot weight (g) | 0.44±0.01<sup>b</sup> | 0.60±0.05<sup>a</sup> | 0.32±0.03<sup>b</sup> | 0.37±0.03<sup>a</sup> | 0.36±0.01<sup>b</sup> | 0.44±0.02<sup>a</sup> |
| Surface (mm<sup>2</sup>) | 47.59±2.88<sup>b</sup> | 54.20±2.51<sup>a</sup> | 36.22±2.32 | 35.50±2.26 | 41.63±1.83<sup>b</sup> | 47.33±2.49<sup>a</sup> |

The simultaneous presence of plant and earthworm resulted in non-neutral reciprocal effects on bacterial communities. Earthworms influence on rhizosphere communities was always detected, (CAP1-2, green and red triangles, Fig.1), with an average abundance increase of earthworm-responding rhizosphere OTUs up to ~3-folds (Supporting File 4). Plant influence on cast communities was also always detected, although weaker in the clayed soil (CAP2, yellow and red squares, Fig.1), with an average abundance increase of plant-responding cast OTUs up to ~9-folds in the sandy soil (Supporting File 4). As cast and rhizosphere community profiles with both macroorganisms can not be predicted from the addition of their coordinates in single macroorganisms treatments (Fig.1), our results demonstrated their significant interaction in shaping microbial communities (Supporting File 3). Nevertheless, the strength of this interaction was modulated by soil type, with a prevalent effect of plant in the sandy soil, and earthworms in the clayed soil.

This plant-earthworm interaction questions the existence of a specific community sub-sample amongst cast and rhizosphere bacteria, selected from *cosmopolitan OTUs present in all soils*. Indeed, we detected a *“core microbiota*”^4,5^ of the earthworm/plant interaction *shared amongst all soils*, featuring 106 OTUs (mostly Alphaproteobacteria and Actinobacteria), always observed in casts and rhizospheres when both macroorganisms are present (purple and red intersect, Fig.2). However, the presence of these ubiquitous OTUs is not necessarily specific to plant-earthworm interaction *per se*. Therefore, we highlighted these community sub-samples using a “source-sink” approach inspired from the metacommunity theory^2,3^ and based on parsimonious hierarchical sorting^17^ to trace back probable origins of OTUs found in cast (g) and rhizosphere (h) microhabitats when both macroorganisms were present (Fig.3a). We observed that the main source of bacteria was the bulk matrix without any macroorganism (a: 72% in g; 65% in h), followed by the microhabitats (bc, 13% in g; 20% in h) and other bulk soils (def, 6% in g; 4% in h). This “source-sink” representation applied to all soils simultaneously revealed again a prevalent contribution of plant (h: 18%, g: 6%) compared to earthworms (g: 7%, h: 2%), and evidenced endemic sub-samples only seen in microhabitats whith both macroorganisms (g: 9%; h: 14%). We propose to call these remaining cast and rhizosphere specific fractions emerging from their simultaneous presence in all soils the “*core microbiota of the plant-earthworm interaction*”.

**Fig.2:**
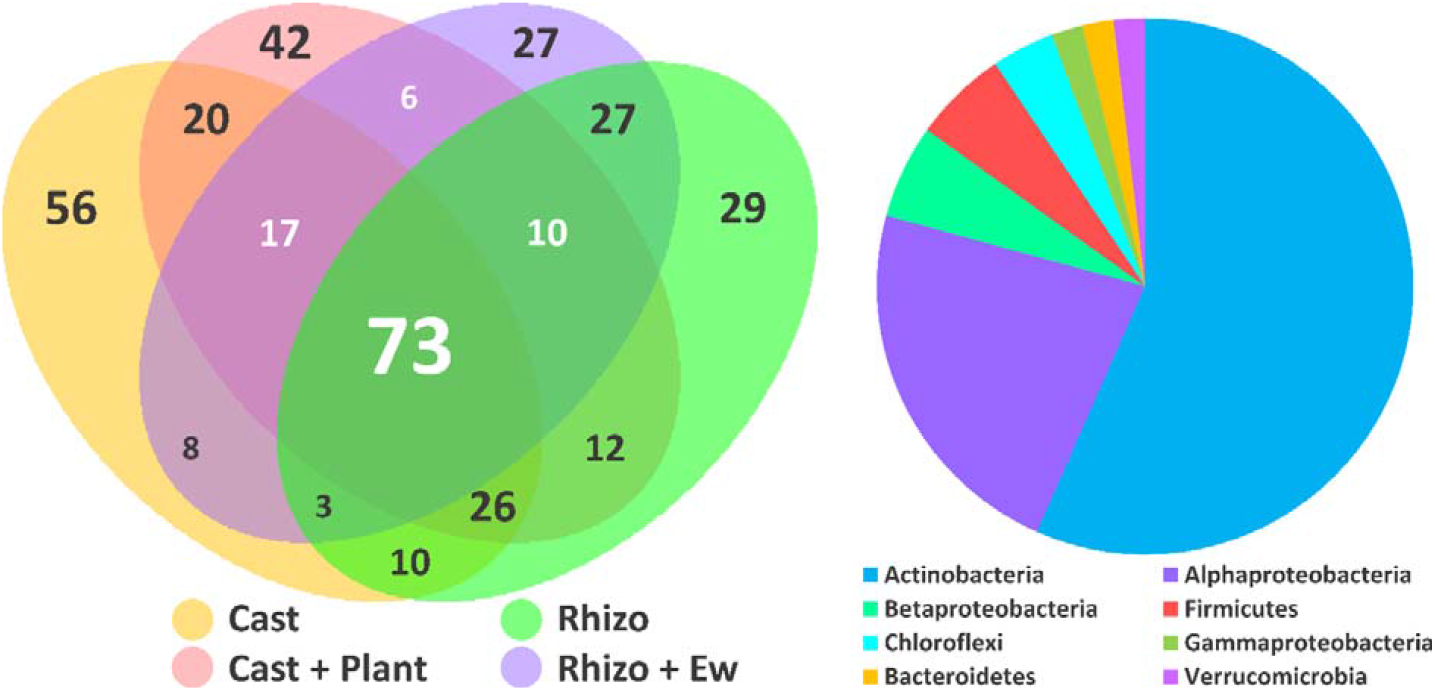
Venn diagram and taxonomy of the core microbiota shared between casts and rhizosphere when both macroorganisms are present (in white, n = 106). Only OTUs found at least in 75% of the biological replicates (3/4) and in all three soils were considered. The piechart shows the unweighted taxonomic distribution of the 106 OTUs, mainly dominated by Actinobacteria (n = 60) and Alphaproteobacteria (n = 24) (n = 48 samples).

**Fig.3:**
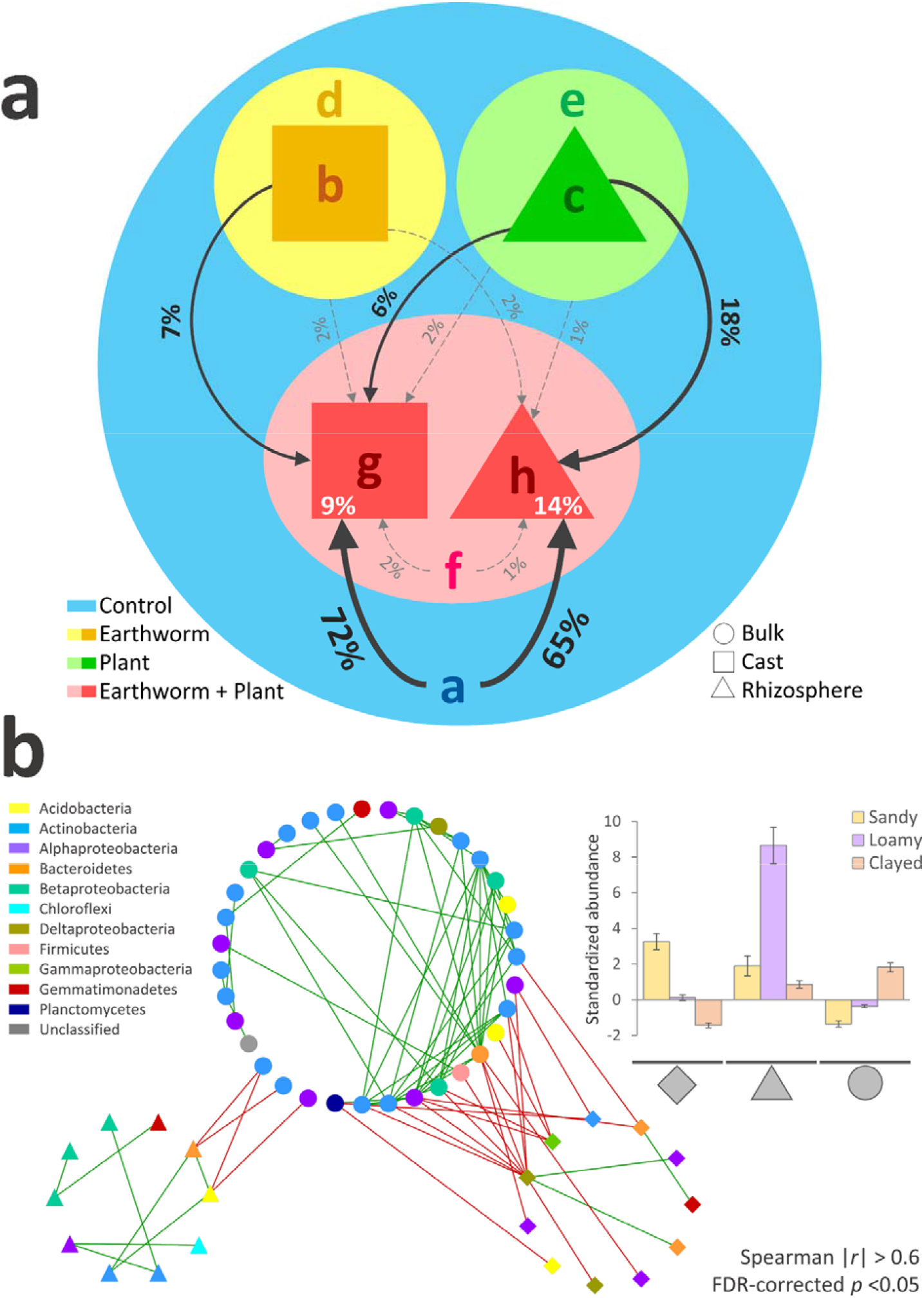
**Panel a.** Source-sink plot tracing the origin of OTUs coming from “sources” (a-f) going into “sinks” (here the cast and rhizosphere when both macroorganisms are present, g and h). Only OTUs found in 75% of biological replicates (3/4) and in the three soils were included (n = 465). Hierachical sorting was applied to attribute OTU source from the following orders respectively for the cast sink “g”: {abchdef} and the rhizosphere sink “h”: {acbgedf}. Source contributions are indicated in percentages, with discontinuous light-grey arrows for contributions < 3%, and no arrows if contribution was null (n = 96). **Panel b.** Core network of OTUs found in cast and rhizosphere of all soils. Module membership (node shapes) was attributed based on OTU standardized abundances against control bulk matrices as shown in the upper-right barchart (average z-score ± standard error of the mean, n = 48). When both macroorganisms were present, diamond-shaped OTUs abundance increased in sandy soil, but decreased in clayed soil. Conversely, circle-shaped OTUs abundance decreased in sandy soils while increasing in clayed soils. Triangle-shaped OTUs increased in all soils, especially in the loamy soil.

The plant-earthworm interaction may also be characterized *via* modifications of OTUs abundance across soils regardless of their origins rather than species composition. We focused on cosmopolitan OTUs found in all soils, and normalized their abundance variations relatively to their control bulk matrix (z-score) in order to build correlation matrices for each microhabitat. We then reconstructed a network specific of the plant-earthworm interaction by keeping only correlations commonly found in cast and rhizosphere matrices, but absent in the bulk (see material and methods). We found a cast and rhizosphere-specific network representing OTUs whose standardized abundance was significantly modulated by plant-earthworm interaction (Fig.3b). Similarly to the Venn diagram (Fig.2), this network was characterized by a phylogenetic signal in favor of Actinobacteria and Alphaproteobacteria. The network was organized in three groups based on hierachical clustering of OTUs z-scores (Supporting File 5). These groups were in line with soil types, indicating differentially altered abundance of these cosmopolitan OTUs when both macroorganisms are present (barplot, Fig.3b). While other studies focused on identification of a *core microbiota* specific to a given host^4,5^, we expanded this concept to the interaction between plant and earthworms *via* OTU correlations, evidencing a *core network* that enables the description of multiple-macroorganisms shaping of bacterial communities.

Moreover, going beyond mere taxonomy, our qPCR results show that the outcome of this interaction may affect other microbial domains. Indeed, the simultaneous presence of earthworms and plants resulted in microbial abundance increase in microhabitats relative to their respective bulk (z-score, Fig.4). This was observed in the loamy and sandy soils for bacteria, but also for fungi in all soils, and archaea in the sandy soil (Fig.4, stars above lines), suggesting that their combined presence have effects going beyond bacteria, impacting the whole microbial community. Moreover, these increase in microbial abudances linked to macroorganism interaction were more frequent in the sandy soil (two bacterial, one archaeal, one fungal, n = 4) compared to the loamy (one fungal, n = 1) and clayed (n = 0) soils, suggesting stronger effects depending on soil type. Additionaly, this interaction was responsible in creation and/or reinforcement of so-called “hotspots”^1^ relative to the bulk soil (Fig.4, stars above bars). Likewise, hotspot numbers were soil-dependent, with higher occurence in the sandy soil (n = 6), compared to the loamy (n = 4) and clayed (n = 2) soils (Fig.4).

**Fig.4:**
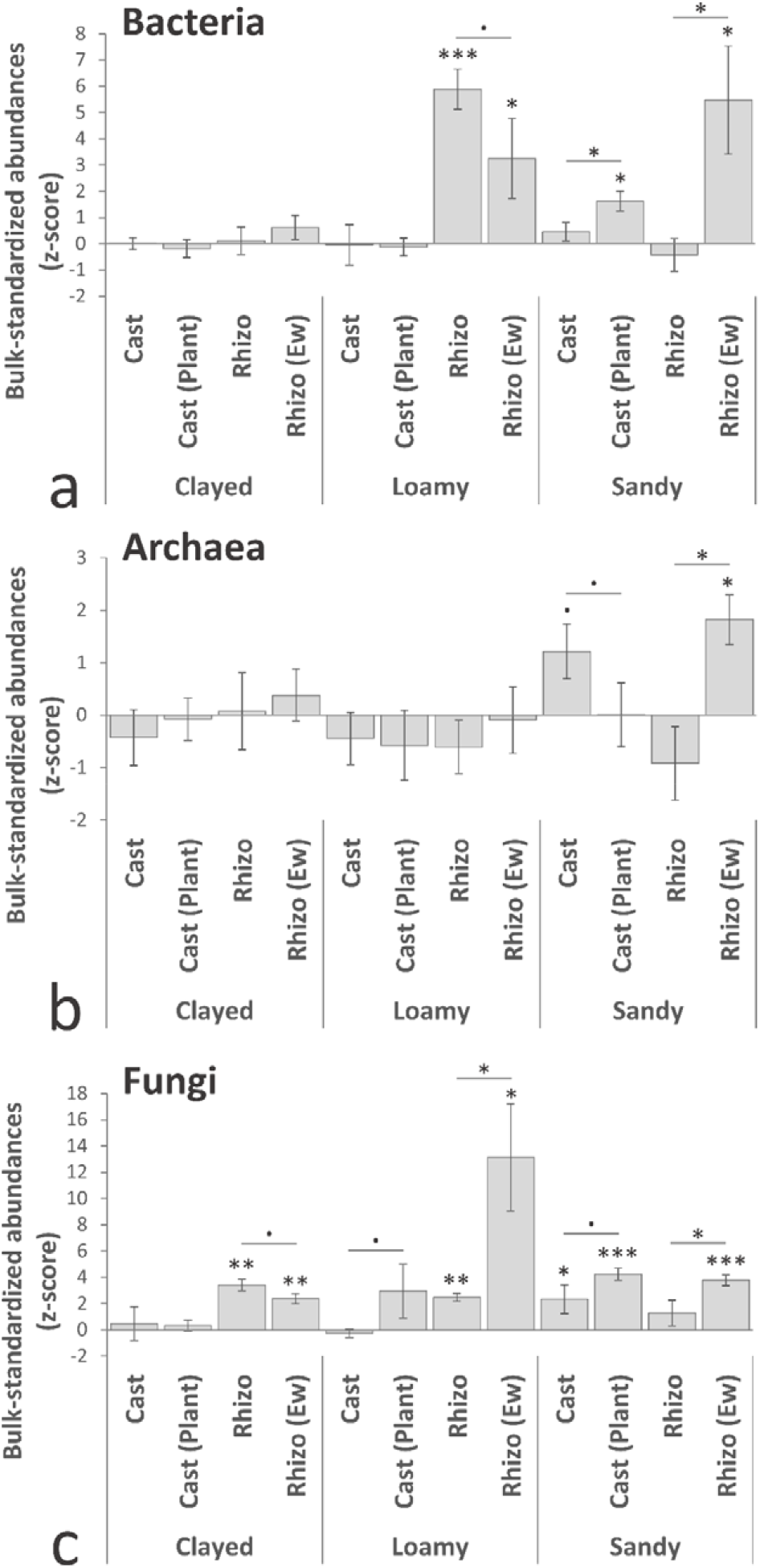
qPCR estimation of bacterial/archaeal genetic markers (a and b, 16S rRNA gene) and fungi (c, ITS). Molecular copy counts were standardized against average and standard deviation values of reference bulk soils from the same treatment (z-score). Barcharts are representing z-score averages ± standard error of the mean (n = 3-5). Significance between treatments were all assessed by two-sample, one-sided Student tests. Significance: *** *p* < 0.001; ** *p* < 0.01; * *p* < 0.05; . *p* > 0.1.

## Conclusion

Altogether, our result indicated a joint shaping of bacterial communities by plant and earthworm, correlating with an increase in plant biomass. This interaction resulted in the emergence of i) *core microbiota* specific of plant-earthworm interaction revealed in cast and rhizosphere while systematically present in all soils, as well as ii) a *core network* commonly shared between cast and rhizosphere whose modularity was indicative of soil type. The contribution of the plant was always dominant. Earthworms influence was soil-dependent, as well as plant-earthworm interaction, whose importance was reinforced in the sandy soil. This joint shaping by two macroorganisms also affected archaea and fungi, especially in the sandy soil. Our data suggest that the impact of the interaction between these macroorganisms on microbial communities is more important when soil is less fertile. Recent studies have accumulated evidences suggesting that microbial community members may be recuited on a functional, rather than a taxonomic basis, alos know as the “It is the song, not the singers” theory^18^. Nevertheless, we show that the *core microbiota* resulting from macroorganism interaction can be highlighted on a phylogenetic basis, with a taxa-specific phylogenetic signal. A perspective based on an adaptive rational would be to investigate these multiple-macroorganisms *core microbiota* and *core network* on a functional basis^19^

## Material and Methods

### i. Microcosm establishment and sampling

Three soils were used: a poor sandy soil classified as cambisoil with moor (organic carbon:14.7 g kg-1; total nitrogen: 1.19 g kg-1; pH: 5.22; clay: 6.9%, loam: 19.0%, sand: 74.1%, origin: CEREEP, Saint-Pierre-Lès-Nemours, France); a loamy crop soil classified as luvisoil (organic carbon: 9.2 g kg-1; total nitrogen: 0.87 g kg-1; pH : 7.0; clay: 16.7%, loam: 56.2%, sand: 27.1%, origin: INRA, Versailles, France) and a forest clayed soil classified as leptosoil (organic carbon: 56.7 g kg-1; total nitrogen: 4.65 g kg-1; pH : 7.45; clay: 34.4%, loam: 39.2%, sand: 27.4%, origin: MNHN, Brunoy, France). Only the first 20cm were sampled, excluding plant debris and roots. Soils were air-dried, sieved (2mm) and set in microcosm pots of 1l containing 1kg of soil watered at 80% of their respective water holding capacity, being the optimum for plant and earthworm^20^. Four conditions were tested with five biological replicates (Supporting File 1): i) plant alone, ii) three earthworms alone, iii) both together and iv) nothing (control). Barley (commercial variety of *Hordeum vulgare* L. from “La fermette”) was germinated in three batches of 80 seeds in Petri dishes containing each soil moisted at 100% (20°C in phytotron, seven days), and ~8cm seedlings were transplanted in pots. The earthworm species *Aporrectodea caliginosa* was chosen for its endogeic lifestyle. Earthworms were coming from a non-stop breeding program started in 2007, with individuals coming from the IRD park (Bondy, France). Three batches of young individuals were purged during three days using their respective experimental soil to avoid massive contaminations from the substrate used for breeding. Three individuals were introduced at the surface of pots, corresponding to a total weight of ~1g. Microcosms were incubated in a climatic chamber (S10H, Conviron, Canada) in the following conditions: 75% air humidity, 18/20°C night/day for a 12h photoperiod at constant light intensity of 300 μmol photons m-2 s-1, for 28 days. Leaf surface was estimated after 17 days by summing leaf areas (one leaf area = leaf lenght x mid-section leaf width x 0.75; leaf-specific correction coefficient for grass-like plants = 0.75)^21^. Plant height was estimated after 23 days based on the length of the longest leaf. Shoot biomass was measured after drying at 50°C for 48h. Soil was meticulously dismantled to sample distinct microhabitats: rhizospheric soil (thightly adhering root soil at 70% humidity, recovered from vigorous shaking with distillated water, then centrifuged), earthworm casts (visual identification)^22^ and bulk soil (remaining soil without visible influence of roots and eathworms).

### ii. DNA extraction and qPCR settings

Total microbial genomic DNA was extracted from 250 mg of soil, collected from the different microhabitats, using FastDNA^®^ SPIN Kit for Soil (MP Biomedicals) following manufacturer’s protocol. DNA concentration was quantified using Quant-iT™ dsDNA High-Sensitivity Assay Kit (Invitrogen) before dilution at 1 ng.μl-1. Abundances of fungal ITS (ITS3F: 5’-GCATCGATGAAGAACGCAGC-3’, ITS4R: 5’-TCCTCCGCTTATTGATATGC-3’)^23^, Crenarchaeota 16S rRNA (Crenar771F: 5’-ACGGTGAGGGATGAAAGCT-3’, Crenar975R: 5’-CGGCGTTGACTCCAATTG-3’)^24^ and bacterial 16S rRNA genes (341F: 5’-CCTACGGGAGGCAGCAG-3', 534R: 5'-CCTACGGGAGGCAGCAG-3')^25^ in samples were achieved using real-time polymerase chain reactions (qPCR) on a StepOnePlus™ Real-Time PCR system (Applied Biosystem, France). The 15μl reaction mixture was composed of 7.5μl Power SYBR™ Green PCR Master Mix with ROX (Applied Biosystem), 1.5μl respective forward and reverse primers (10μM), 2.5μl UltraPure™ DNase/RNase-Free Distilled Water (Applied Biosystem) and 2μl DNA template. Potential qPCR reaction inhibition by DNA matrix was assessed by doing a preliminary analysis adding 2μl of known concentration of plasmid to reaction mixture previously described thus adjusting water volume to 0.5μl. No inhibition was detected as amplification using primers targeting T7 and SP6 RNA polymerase promoters was similar among samples. Conditions for real-time PCR were 900 s at 95 °C for enzyme activation followed by 35 cycles of 15 s at 95 °C, 30 s at specific annealing temperature (ITS/Archaea: 55°C, Bacteria: 60°C), 30 s at 72 °C for elongation and 30 s at 80 °C for data collection. Each abundance was quantified with three repetitions using linearized plasmid-based standard curve with StepOne™ Software v2.2.2. Deviation between the different qPCR plates was corrected using samples calibrator presents in each plate. Amplicon copy numbers were normalized per μgram of DNA per gram of soil, transformed with log2, and used to build linear models. Normality of the data was assessed with d’Agostino test on the residuals of each model (bacteria *p* = 0.17, fungi *p* = 0.72, archaea *p* = 0.36). Outlier values were removed based on ANOVA diagnosis plots. Respectively 3/96, 4/96 and 3/96 values were removed for bacteria, fungi and archaea respectively, leaving between 3-5 biological replicate values per condition. To account for the strong soil and macroorganism effects, rhizopshere and cast datasets were standardized using their respective bulk soils under the macroorganisms presence (z-score). Statistical significance against the bulk soil of the control treatment and between microhabitats was tested with Student tests (one-sided, two-sample, *p* < 0.05). The one-sided version of the test was selected as we hypothesized that macroorganisms will have a positive impact on molecular abundances of soil microorganisms.

### iii. 16S rRNA gene amplicon sequencing and bioinformatics

Amplicons were generated from purified DNA in TE buffer by LGC Genomics (GmbH, Germany), respecting the best practices guidelines^26,27^. In the first step, the bacterial 16S rRNA gene V3-V4 hypervariable region was PCR-amplified using the fusion primers U341F (5’-CCTACGGGNGGCWGCAG-3’) and 785R (5’-GACTACHVGGGTATCTAAKCC-3’)^28^. For each sample, the forward and reverse primers had the same 10-nt barcode sequence. PCR was carried out in 20μL reactions containing containing 1.5 units MyTaq DNA polymerase (Bioline, Germany) and 2 μl of BioStabII PCR Enhancer (Sigma, Germany), 15pmol of each primer, and 5 ng template DNA. Thermal cycling conditions were 96°C for 2 min followed by 30 cycles of 96°C for 15s, 50°C for 30sec and 70°C for 90s, with a final extension at 72°C for 10 min. PCR products were visualized in 2% agarose gel to verify amplification and size of amplicons (around 500 bp). About 20 ng amplicon DNA of each sample were pooled for up to 48 samples carrying different barcodes, thus two pools of 48 samples were generated (n = 96). The amplicon pools were purified using AMPure XP beads (Agencourt, Germany), followed by an additional purification on MinElute columns (Qiagen, Germany). About 100 ng of each purified amplicon DNA pool was used for Illumina library construction using the Ovation Rapid DR Multiplex System 1-96 (NuGEN, Germany). Illumina libraries were then pooled and size selected by preparative Gel electrophoresis. Sequencing was performed on MiSeq (Illumina, 2×250 bp) using the MiSeq reagent kit v2. Demultiplexing and trimming of Illumina adaptors and barcodes was done with Illumina MiSeq Reporter software (version 2.5.1.3). Sequence data were analyzed using an in-house developed Python notebook piping together different bioinformatics tools (available upon request). Briefly, sequences were assembled using PEAR^29^ with default settings, removing short sequences and quality checks of the QIIME pipeline^30^. Reference based and *de novo* chimera detection, as well as clustering in OTUs were performed using VSEARCH^31^ and the Greengenes reference database. The identity thresholds were set at 97%. Representative sequences for each OTU were aligned using PyNAST^32^ and a 16S phylogenetic tree was constructed using FastTree^33^. Taxonomy was assigned using UCLUST^34^ with the latest released Greengenes database^35^, and the final contingency table was set at OTU level. Rarefaction curves were calculated with the *vegan* package^36^ in Rgui^37^ to assess sequencing depth and samples were rarefied to 6900 counts.

### iv. Beta-diversity and multivariate analysis

Rarefied OTU matrices and unifrac trees were used to build variance-adjusted weighted and unweighted unifrac-based constrained analysis of principal coordinates (CAP, *capscale* function, package *vegan*). Models were validated with 10,000 permutations. Responding OTUs whose abundance was significantly altered in rhizosphere and casts when both macroorganisms were present (but not in their respective bulk) were extracted using quasi-likelihood F-test under negative binomial distributions and generalized linear models (nbGLM QLFT, FDR-adjusted *q* < 0.05). OTUs significantly affected by the addition of a second macroorganisms in microhabitats were extracted as previously described^38^ *via* hierarchical clustering in heatmaps for cast (Supporting File 6) and rhizosphere samples (Supporting File 7), followed by grouping in barcharts (Supporting File 5).

### v. Venn diagram and source-sink plot

A Venn diagram was done with the Rgui package *limma*^39^ to define the core microbiota shared between cast and rhizosphere using only cosmopolitan OTUs strictly found at least in 75% of biological replicates (3/4) in all three soils (Figure 2). A “source-sink” plot was established to trace OTU’s origin, using cosmopolitan OTUs found in 75% of biological replicates (3/4) in all three soils. (Fig.2). We hypothesized that the sources of bacteria for casts (g) and rhizosphere (h) communities in the presence of both macroorganisms were as follow: 1) the initial soil matrix without macroorganisms (a); 2) the microhabitat created by each macroorganism alone (c for g and e for h); 3) the second microhabitat created by the other macroorganism (e for g and c for h); 4) the other microhabitat when both macroorganisms were present (h for g and g for h); 5) the bulk soil surrounding the microhabitats from macroorganisms alone (b for g and d for h); 6) the bulk soil surrounding the other microhabitats from macroorganisms alone (d for g and b for h); 7) the bulk soil when both macroorganisms were present (f); 8) the remaining part was specifically attributed to each microhabitat under interaction context as endemic fractions (g and h).

### vi. Core microbial network

To account for the strong soil effect, OTU abundances in each sample were standardized to their respective control bulk average and standard deviation without macroorganisms (z-score). We focused on cosmopolitan OTUs present at least in 50% of samples in each soil (n > 16/32) and used their standardized abundances to build three correlation networks, one per microhabitat, using stringent cut-off (Spearman’s rho < |0.6|, FDR-adjusted *q* < 0.05). This resulted in a “bulk network” (originating from all the standardized bulk samples in earthworm, plant, earthworm/plant treatments in the three soils, n = 36), a “cast network” (originating from all the standardized cast samples in the three soils, n = 24), and a “rhizosphere network” (originating from all the standardized rhizosphere samples in the three soils, n = 24). Hereafter, we intersected the cast and rhizosphere networks to only keep the overlapping correlations (exclusions and co-occurrences) in common between these two microhabitats. Last, we removed any bulk interference correlations from the rhizosphere-cast intersected network by subtracting correlations from the bulk network. This network arithmetic was done with the Rgui package *igraph*^40^. Modularity in the network was attributed based on hierarchical clustering of their standardized abundances (Supporting File 5), which was summarized in the upper-right barplot of Fig. 3, classifying OTUs in three modules based on their average z-score behavior across the three soils.

## Acknowledgements

We thank Valérie Serve for technical help, Beatriz Decencière, Amandine Hansart and Florent Massol of the CEREEP - Ecotron IDF/UMS CNRS/ENS 3194 for the sandy soil, Sandrine Salmon of the UMR 7179 / CNRS-MNHN for the clayed soil and Christophe Montagnier of the UE Grandes cultures / INRA for the loamy soil. This work was supported by grants from the French national program CNRS/INSU [EC2CO-Biohefect-MicrobiEn-AuxAzote].

## Author contributions and information

SJ (analytical strategy, data analysis, manuscript writing), RPF (laboratory experiment, manuscript editing), CMon (sequencing strategy), CMou (sequencing strategy), AS (analytical strategy, data analysis, manuscript editing), AM (bioinformatic), LP (analytical strategy, manuscript editing), MB (study conception, research direction, analytical strategy, manuscript writing). Authors declare no competing interests.

## Code and data availaibility

The Rgui software and associated function packages used for data analysis are all publically available. Data that support the findings of this study have been deposited in the Sequence Read Archive database (SRA, https://www.ncbi.nlm.nih.gov/sra) with the primary accession code “SUB5123378”, and will be made automatically publically available after publication.

