## Supporting Files for "A core microbiota of the plant-earthworm interaction conserved across soils"

–

**Supporting File Legends**

**Supporting File 1:** Experimental design. Three soils (sandy, loamy, clayed) were crossed with four treatments (no macroorganisms, earthworms, plant, both) and replicated five times, giving 60 microcosms. Plant traits were recorded on each microcosms (n = 60). Soil microhabitats were sampled depending on the treatment, yielding 40 samples per soil type (20 bulks, 10 casts and 10 rhizospheres), representing a total of 120 samples (3 soils x 40 microhabitats). qPCR assay was done on all samples (n = 120) while 16S rRNA gene amplicon sequencing was done only on four biological replicates, excluding the outlier-most sample based on plant traits (n = 96).

**Supporting File 2:** Canonical analysis of principal coordinates of bacterial communities based on variance-adjusted weighted unifrac distances on the three soils together. CAP1 and 2 represent the constrained components with their respective percentage of the total variance explained. The model was validated using 10.000 permutations. The soil was the main structuring factor (colors) separating on the first two components (50.6%), while the differences related to microhabitats (dot shape) were nested inside each soil (n = 96).

**Supporting File 3:** Nested analysis of variance (ANOVA) of bacterial communities exposed to plants, earthworms, or both (weighted unifrac distance matrix, 10.000 permutations, adjusted-$r^2$ = 0.7***, $p$ < 1.0E-6). The model tested was constructed to nest microhabitat creation within macroorganisms presence. Soil (clayed, loamy, sandy) and microhabitats (bulk, cast, rhizosphere) were set as categorical factors. Macroorganism factor (earthworm and plant) was set as a continuous variable (1 = presence, 0 = absence) (n = 96).

**Supporting File 4:** Groups of OTUs reacting to the second macroorganisms in casts and rhizospheres (see Supporting File 6 and 7 for details on OTUs selection). An increase of 2.6, 2.2 and 2.9 folds as well as a decrease of 1.5, 2 and 4.7 folds were observed for responding OTUs respectively in the sandy, loamy and clayed rhizospheres with earthworms (panels ac).

Likewise, an increase of 8.9, 4.2 and 1.8 folds as well as a decrease of 3.5, 1.6 and 1.6 folds was observed for responding OTUs respectively in the sandy, loamy and clayed casts with plants (panels bd) (n = 96).

**Supporting File 5:** Membership of OTUs found in the rhizosphere-cast network interaction based on their standardized abundance against their own bulk soil matrix (z-score). The origin of samples (soil and microhabitat) is indicayed by the colored symbols in the top dendrogram. The OTU profiles were clustered on the left dendrogram into three groups based on their standadrized abundance (z-score, diamond = increase in the sandy soil, triangle = increase in the loamy soil rhizosphere, circle = increase in the clay and loamy soils).

**Supporting File 6:** Cast OTUs significantly responding to plants in the three soils (Quasi-Likelihood Ratio tests, generalized linear model with negative binomial distribution, FDR-adjusted $q < 0.05$). Samples information is presented on the leftside color key (rows). OTU abundance was centered and scaled by columns for visualisation (z-score), and clustered (method = “complete”, distance = “Euclidean”). Each heatmap represents cast OTUs either significantly increased (green plus) or decreased (red minus) by plants (purple) relative to their intital abundance in the control bulk (blue), the bulk soil with both macroorganisms (pink), and the cast without plants (yellow).

**Supporting File 7:** Rhizosphere OTUs significantly responding to earthworms in the three soils (Quasi-Likelihood Ratio tests, generalized linear model with negative binomial distribution, FDR-adjusted $q < 0.05$). Samples information is presented on the leftside color key (rows). OTU abundance was centered and scaled by columns for visualisation (z-score), and clustered (method = “complete”, distance = “Euclidean”). Each heatmap represents rhizosphere OTUs either significantly increased (green plus) or decreased (red minus) by earthworms (purple) relative to their intital abundance in the control bulk (blue), the bulk soil with both macroorganisms (pink), and the rhizosphere without earthworms (green).

**Supporting File 1**

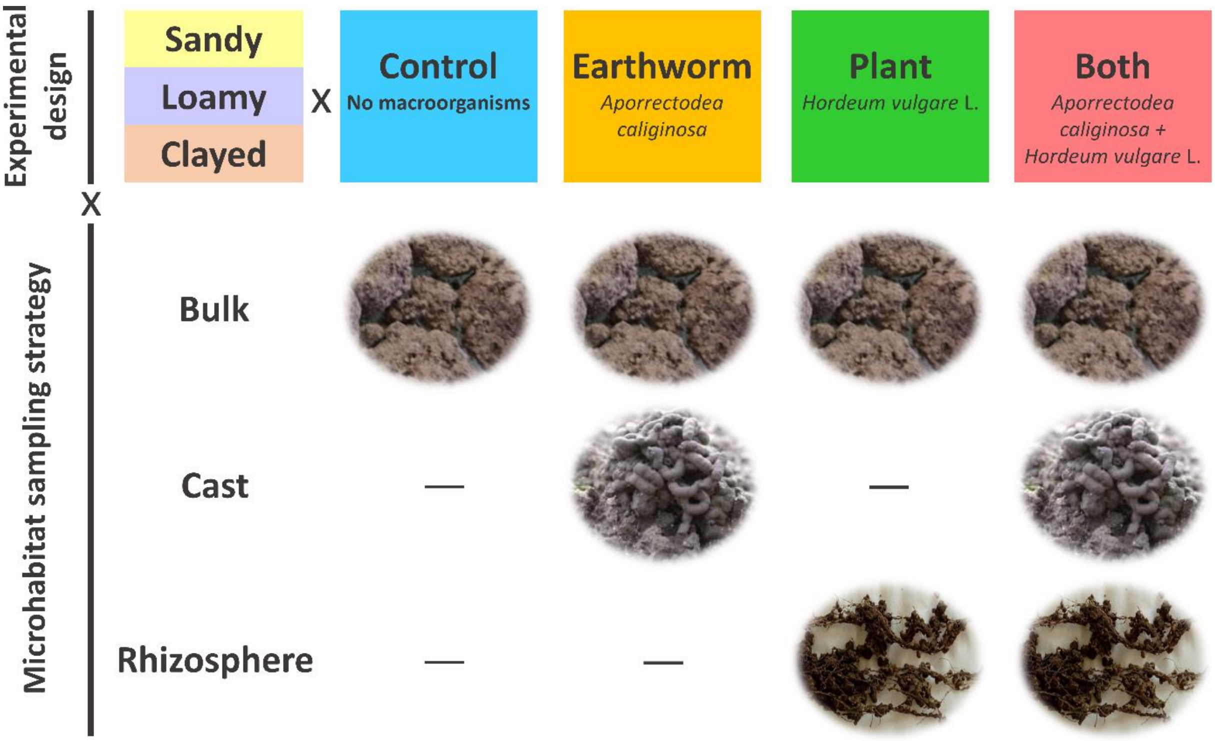

**Supporting File 2**

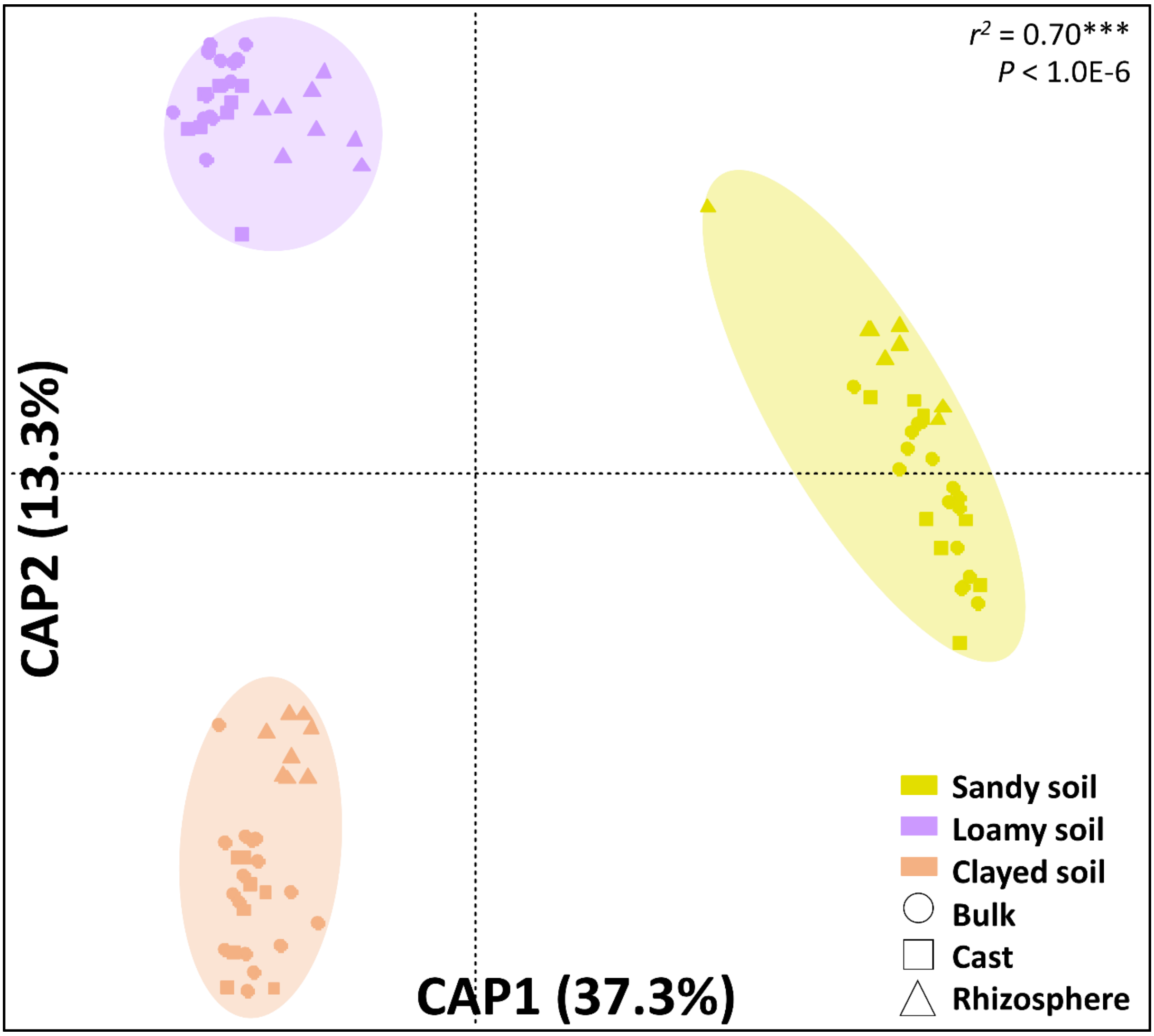

**Supporting File 3**

| Factors and interactions tested | Var. (%) | *p* | Signif. |
|---|---|---|---|
| Soil | 51.79 | 1.00E-6 | *** |
| Plant | 3.91 | 1.00E-6 | *** |
| Earthworm | 0.83 | 0.05 | * |
| Soil:Plant | 2.04 | 6.90E-3 | ** |
| Soil:Earthworm | 1.28 | 0.08 | . |
| Plant:Earthworm | 0.68 | 0.09 | . |
| Soil:Plant:Earthworm | 0.87 | 0.30 | - |
| Plant:Earthworm:Microhabitat | 4.75 | 1.00E-6 | *** |
| Soil:Plant:Earthworm:Microhabitat | 3.65 | 1.30E-3 | ** |
| Residual | 30.2 | - | - |

**Supporting File 4**

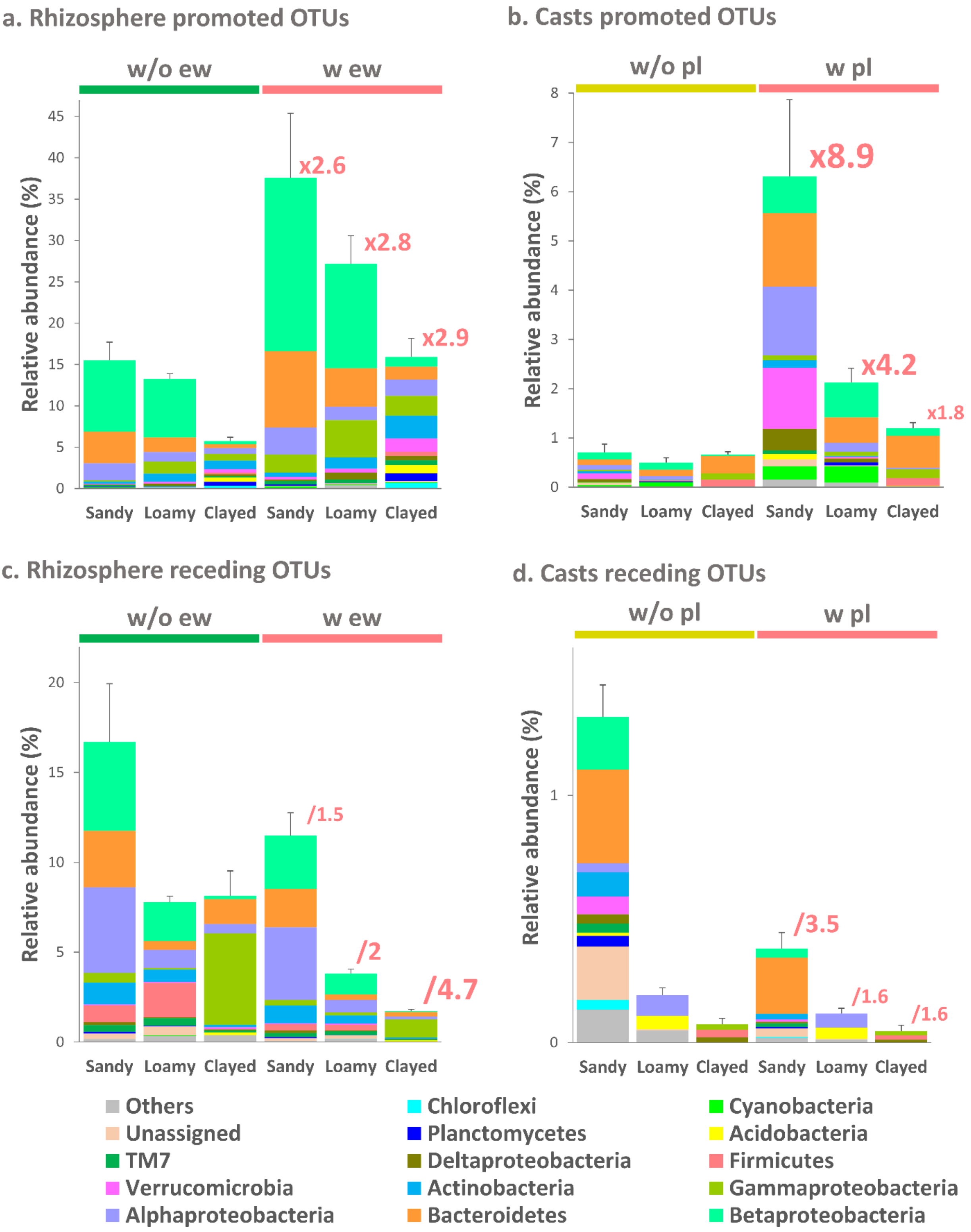

**Supporting File 5**

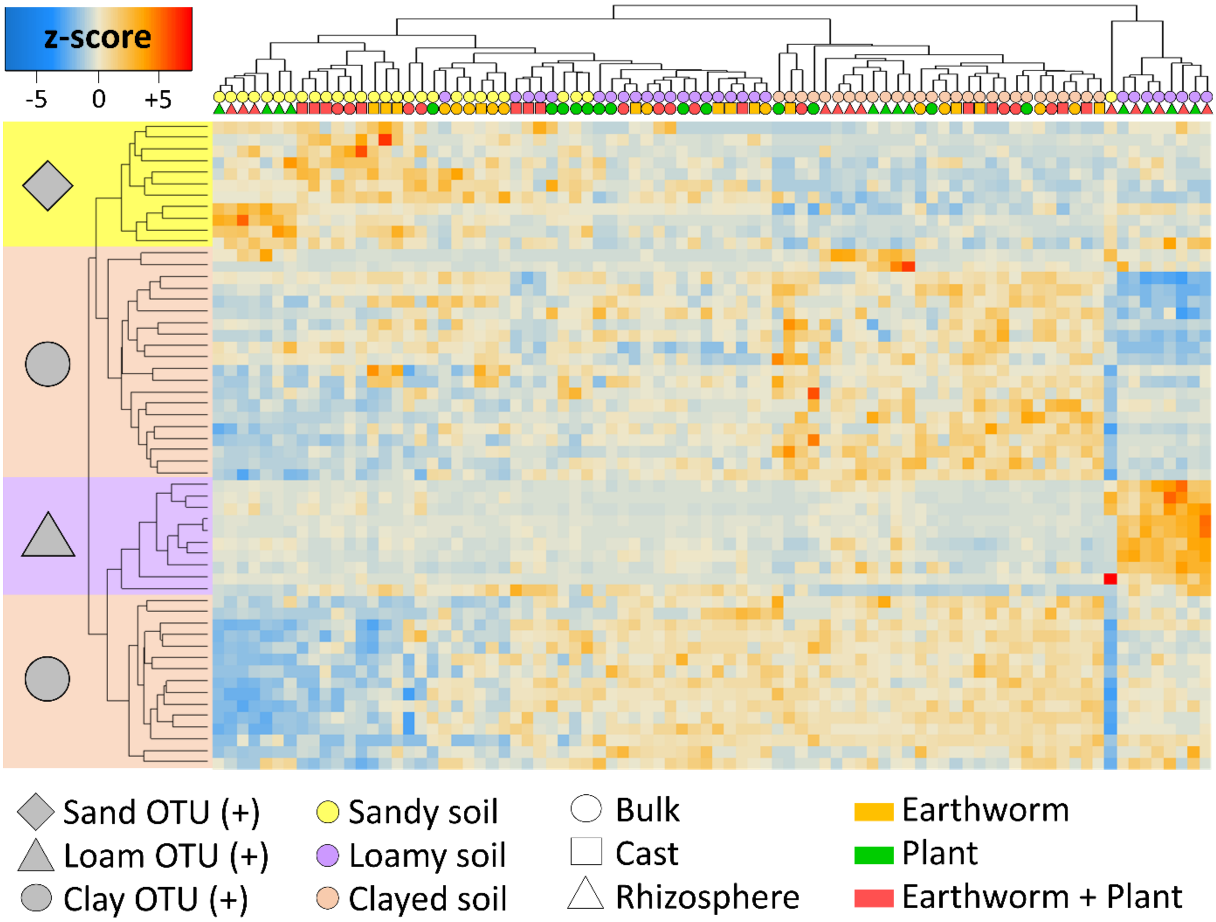

**Supporting File 6**

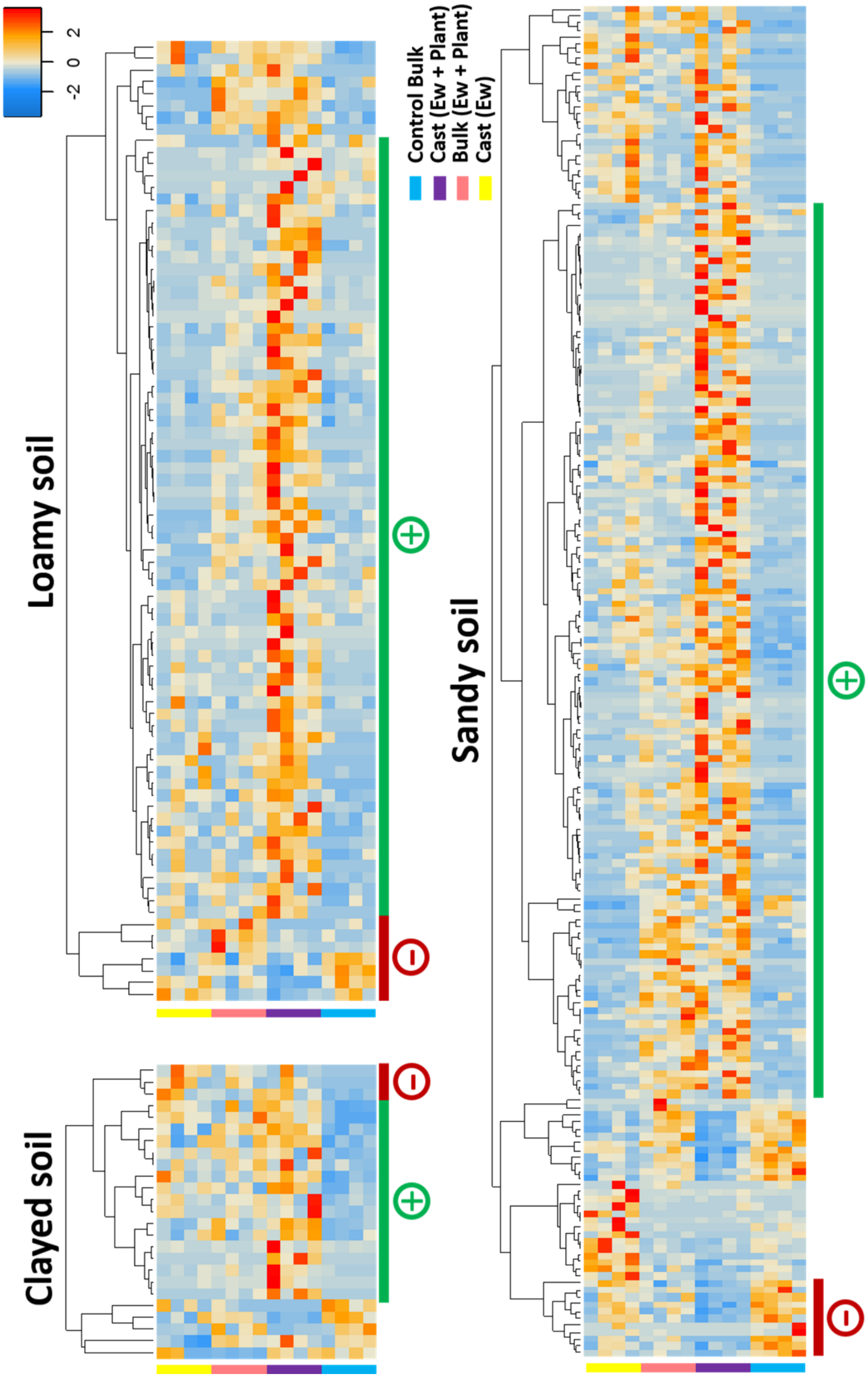

**Supporting File 7**

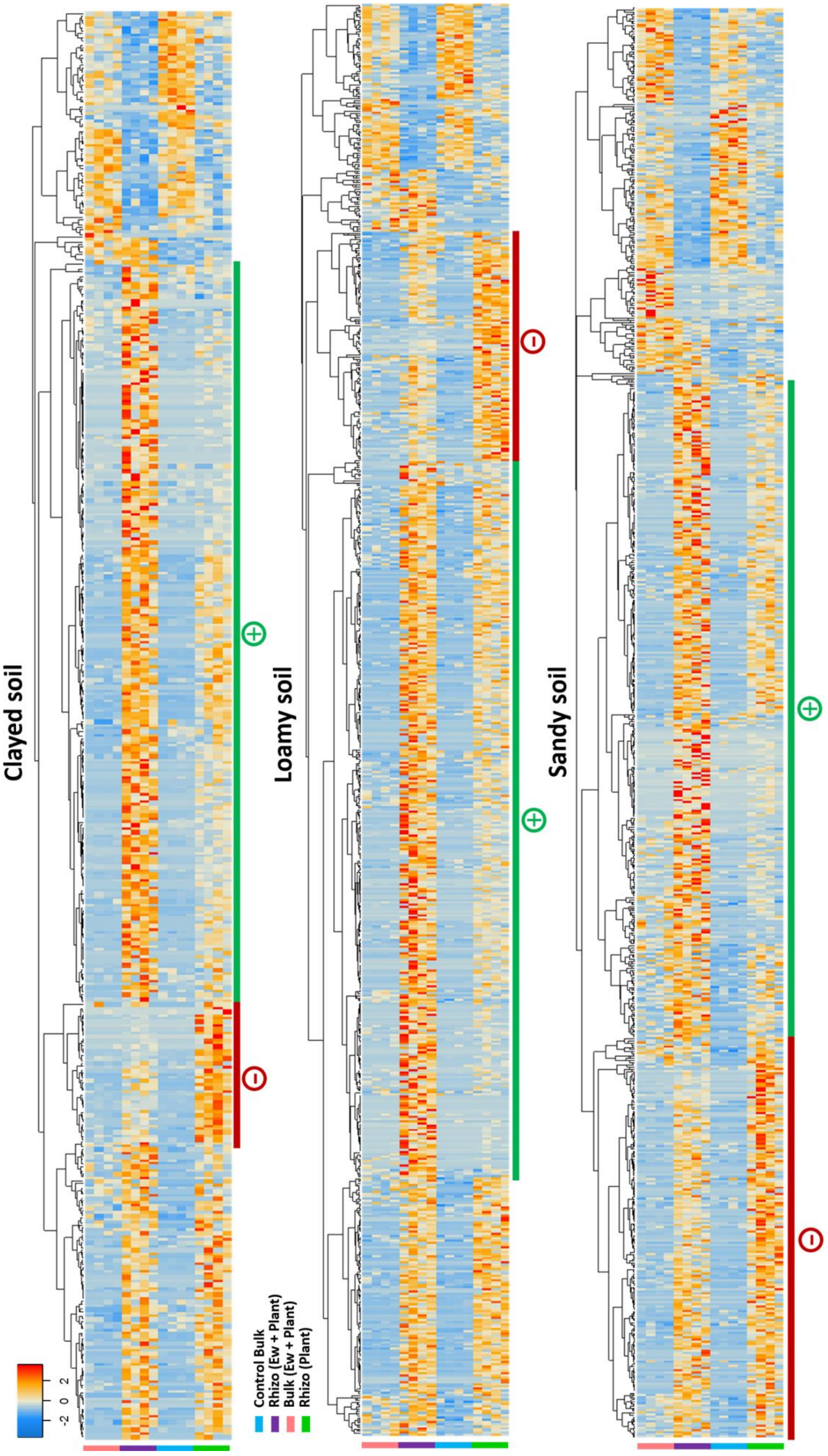
